# Gamete release in *Ciona robusta*: roles of gonadotropin-releasing hormone and the photoreception system

**DOI:** 10.1101/2025.07.01.661410

**Authors:** Tomohiro Osugi, Shin Matsubara, Akira Shiraishi, Azumi Wada, Yuki Miyamoto, Issei S. Shimada, Yasunori Sasakura, Takehiro G. Kusakabe, Honoo Satake

## Abstract

Gamete release, an essential event for animals, is regulated by various factors including environmental stimuli, neural circuit, and endocrine molecules. In this study, we investigated the mechanisms regulating gamete release in the ascidian *Ciona robusta*, part of the sister group of vertebrates. *Ciona* is a hermaphrodite, releasing sperm first from orange-pigmented organ (OPO) at the end of the spermiduct, followed by eggs from the oviduct beneath the spermiduct knob-like structure. In this study, behavioral and morphological analyses revealed that this sequential release occurs because the knob-like structure compresses the oviduct opening at the onset of gamete release. Observations of transgenic *Ciona* expressing Kaede under the gonadotropin-releasing hormone 2 (*Gnrh2*) promoter showed that *Gnrh2*-expressing neurons and fibers accumulate in the epithelium around the spermiduct openings. In contrast, *Gnrh1*-expressing neurons are localized in the cerebral ganglion and project toward the ovary, suggesting distinct roles of GnRH1 and GnRH2 in reproductive regulation. RNA-seq and real-time PCR analyses revealed that *Opsin2, Opsin3*, beta-carotene-15,15’-monooxygenase (*Bco*), and several ion channel genes are specifically expressed in the OPO, along with *Gnrh2. In situ* hybridization showed that these genes are localized in the innermost OPO layer, suggesting that the OPO functions as a photoreceptive organ. Although, *Gnrh2* expression has been considered low in adult *Ciona*, our study revealed strong OPO-specific expression with photoreceptive genes. Collectively, these findings suggest that GnRH2 plays a central role in gamete release regulation, potentially in coordination with the photoreceptive system, while GnRH1 may regulate ovarian functions. These results indicate that *C. robusta* employs distinct GnRH systems in a tissue-specific manner to regulate reproduction.

## Introduction

Gamete release, an essential event for animals, is regulated by environmental stimuli and various biological factors including neural circuit and endocrine molecules. The mode and timing of gamete release is important especially for aquatic animals because the site of fertilization is in water. Light is one of the common environmental stimuli inducing gamete release. In jelly fish, light directly stimulates gonadal photosensory-neurosecretory cells which induce gamete maturation and release.^1^ In fish, the timing of spawning is tightly connected with the lunar and diurnal cycles during the spawning season in pufferfish and the molecular mechanisms regulating spawning are well established.^2-5^ In ascidians, the sister group of vertebrates, the relationship between light and gamete release was previously studied in *Ciona intestinalis* while interestingly, the timing of spawning is still controversial. Several studies reported that spawning generally takes place at sunrise.^6-8^ On the other hand, it was reported that spawning took place during the evening and night-time hours.^9^ Furthermore, the other study reported that *Ciona* may spawn and settle at any time of the day.^10^ In contrast to the controversy, continuous illumination prevents uncontrolled spawning,^11,12^ suggesting that light is an essential factor inhibiting spawning behavior in *Ciona*.

*Ciona* is a hermaphroditic animal that releases both sperm and eggs. Sperm is released first from the distal end of spermi duct, where orange-colored cells accumulate to form a structure known as orange-pigmented organ (OPO).^13^ Egg release occurs from the oviduct beneath the spermiduct knob-like structure at the OPO after a period of sperm release. Interestingly, orange-colored cells are typically observed in the OPO of *Ciona intestinalis* type A, which is distributed in the Mediterranean, northeast Atlantic, and Pacific regions, whereas in *Ciona intestinalis* type B, found in the north Atlantic region, pigmentation is restricted to the spermiduct.^14^ Based on geographical distribution, morphological characteristics, and genetic information, *Ciona intestinalis* is now be reclassified into two distinct species: *Ciona robusta* (formerly *Ciona intestinalis* type B) and *Ciona intestinalis* (formerly *Ciona intestinalis* type A).^14,15^

Neuropeptides are important regulators of reproduction in animals. Among them, gonadotropin-releasing hormone (GnRH) plays a central role in the hypothalamus-pituitary-gonadal (HPG) axis in vertebrates. In tetrapods, GnRH is generally believed to promote gonadal maturation by stimulating the secretion of pituitary gonadotropins: follicle-stimulating hormone (FSH) and luteinizing hormone (LH).^16-18^ In fish, GnRH has also been reported to regulate spawning behavior.^19,20^ More recently, a “dual GnRH model” has been proposed in fish for the regulation of FSH and LH secretion, in which GnRH stimulates LH secretion, while cholecystokinin stimulates FSH secretion.^21^ The origin of GnRH can be traced back to a common ancestor of bilaterians, forming a superfamily that includes GnRH, adipokinetic hormone, and corazonin, and acts as a multifunctional peptide in both protostomes and deuterostomes.^22-24^ In the deuterostome lineage, GnRH genes have been well conserved. In *Ciona*, two GnRH genes (*Gnrh1* and *Gnrh2*) and their encoded GnRH peptides, and four GnRH receptor genes have been identified.^25-31^ These GnRH and receptor genes are thought to have emerged by lineage-specific duplication events.^32^ GnRH plays essential roles during the development of *Ciona*, which are independent of the regulation of reproduction.^33-35^ In adult *Ciona*, morphological analyses have shown that GnRH immunoreactive neurons and nerve fibers directly reach the ovary, suggesting that GnRH is involved in reproductive regulation.^36^ Interestingly, GnRH injection into gonoduct induced gamete release, while the molecular mechanisms underlying this process remain to be clarified.^26,37^ In addition, the relationship between the GnRH system and above-mentioned light/dark conditions that induce gamete release remains enigmatic.

In this study, we focused on *Ciona robusta* (hereafter referred to as *C. robusta*) and investigated the mechanisms regulating gamete release. Behavioral observations and histological analyses revealed that the sequential release of sperm and eggs occurs in this order because the knob-like structure on the spermiduct physically compresses the oviduct opening during the initial stage of gamete release. This structural characteristic suggests that *C. robusta* actively regulates sperm release but not egg release. Anatomical observations of transgenic *C. robusta* expressing Kaede under the control of the *Gnrh1* or *Gnrh2* promoter further revealed distinct morphological differences between GnRH1 and GnRH2 nervous systems: GnRH1 neurons in the cerebral ganglion project to the ovary, whereas GnRH2 neurons in the OPO project to the cerebral ganglion. Gene expression and histological analyses of the OPO further revealed that not only *Gnrh2* but also photoreception-related genes, including *Opsin2, Opsin3*, and *beta-carotene 15,15’-monooxygenase* (*Bco*), and several ion channel genes are specifically expressed in the OPO. These findings suggest that GnRH2 and the photoreceptive system may represent a missing link in the regulation of gamete release in *C. robusta*.

## Materials and Methods

### Animals

Wild adult *C. robusta* was used for behavioral and morphological observations, gene expression analyses, and histochemical studies. *C. robusta* was also used to generate transgenic lines. The CiGnRH1>*Kaede* lines and CiGnRH2*>Kaede* lines of *C. robusta*, Tg[MiCignrh1K]2, Tg[MiCignrh2K]1 and Tg[MiCignrh2K]2^38,39^, which were created by Minos transposon-mediated transgenesis, were used for anatomical observations of the GnRH1 and GnRH2 nervous systems. No significant differences were observed between the Tg[MiCignrh2K]1 and Tg[MiCignrh2K]2 lines. The transgenic lines were cultured using an inland culture system as described previously^40^.

### Observation of spawning behavior

Wild *C. robusta* were placed in a 30-cm cubic tank and maintained under dark conditions for 30 minutes to 1 hour by covering the tank with a box draped with a blackout curtain. Spawning behavior was monitored under the dark conditions or after removing the box, and video recordings were captured using a digital camera (PENTAX K-70; RICOH IMAGING COMPANY, LTD., Tokyo, Japan), and an action camera (GoPro HERO13 Black; GoPro, Inc., San Mateo, CA, USA) with close-up lens (AOI UCL-03; Fisheye Co.,Ltd., Tokyo, Japan). For observations under a stereomicroscope, wild *C. robusta* were anesthetized using L-menthol according to a previously reported method^41,42^. Briefly, a 0.56% (weight/volume) L-menthol solution in ethanol was prepared and used as a stock solution. The stock solution was further diluted with 1% (volume/volume) artificial seawater before use. Animals were soaked in the diluted L-menthol solution for 10 minutes. After anesthetization, parts of the tunic, and body-wall muscle were excised, and the OPO was exposure under a stereomicroscope (ZEISS SteREO Discovery.V8; Carl Zeiss, Oberkochen, Germany). Gamete release was observed, and video recordings were captured using the stereomicroscope equipped with a USB digital camera (3R-DKMC04; 3R SOLUTION CORP., Fukuoka, Japan) and Anyty microscope adapter software ver. 9.0 (3R SOLUTION CORP.).

### Whole-mount tissue preparation

Transgenic animals were anesthetized as described above. Parts of the tunic, body-wall muscle, and pharynx were excised under a fluorescence stereomicroscope (Leica M205 FA; Leica Microsystems, Wetzlar, Germany), and whole animals were fixed in 4% paraformaldehyde in phosphate buffer (PB) at 4 °C overnight. The fixed specimens were then soaked in 2 mg/ml glycine in PBS to quench the paraformaldehyde. Subsequently, the specimens were washed three times with PBS and used for morphological analyses. Anatomical analyses were performed under the stereomicroscope (Leica M205 FA; Leica Microsystems) using precision tweezers (Outils Rubis SA, Stabio, Switzerland) and microscissors (Inami & Co., Ltd., Tokyo, Japan).

### Whole-mount observation

After removal of the tunic, fixed tissues were placed either in a glass dish with a thick silicone resin sheet or in a slide glass chamber (AGC TECHNO GLASS, Tokyo, Japan) filled with PBS and secured with needles as appropriate. The fluorescence stereomicroscope (Leica M205 FA; Leica Microsystems) and image acquisition software (Leica AF6000E; Leica Microsystems) were used for low-magnification observations.

### Histological analysis of tissue sections of the OPO

The OPO tissues were fixed with Bouin’s fixative solution at 4 °C overnight. The fixed tissues were then soaked in 30% sucrose in PBS at 4^°^ C overnight. The tissues were embedded in Tissue-Tek Cryomold #4565 (Sakura Finetek Japan Co., Ltd., Tokyo, Japan) using Super Cryoembedding Medium-L1 (Leica Microsystems Japan, Tokyo, Japan). The tissues were subsequently frozen in n□hexane cooled to ™196 °C with liquid nitrogen and stored at ™80 °C until sectioning. Sections of 10 μm thickness were prepared using a CryoStar NX70 cryostat (Thermo Fisher Scientific, Inc., Waltham, MA) at ™20 °C. Hematoxylin–eosin staining was performed following a modified version of a previously reported method^42^. Briefly, the sections were washed with distilled water for 5 minutes and immersed in Mayer’s hemalum solution for 10 seconds. They were then washed with PBS for 2 minutes and distilled water for 1 minutes, followed by immersed in eosin solution for 2 minutes. After rinsing with distilled water, the sections were dehydrated through a graded ethanol series (70%, 80%, 95%, and 100%) and xylene. Finally, the sections were mounted with a cover glass (Matsunami Glass Ind., Ltd.) and MOUNT-QUICK (Daido Sangyo Co., Ltd., Tokyo, Japan), and observed under a light microscope equipped with 20× or 40× objectives (Axio Imager 2; Carl Zeiss, Oberkochen, Germany) in phase-contrast mode. OPO tissues of Tg[MiCignrh1K]2 line were fixed in 4% PFA and sections of 10 μm thickness were prepared as described above. Sections were washed three times with PBS for 5 minutes and mounted with a cover glass (Matsunami Glass Ind., Ltd.) and Fluoromount (Diagnostic BioSystems, Pleasanton, CA, USA). The sections were observed under a fluorescent microscope with with 20× or 40× objectives (Axio Imager 2; Carl Zeiss), or a confocal microscope equipped with 100× objectives (FLUOVIEW FV3000; Olympus, Tokyo, Japan).

### RNA-sequence analysis

Animals were anesthetized as described above. OPO, spermiduct, oviduct, and ovary were dissected under a stereo microscope (ZEISS SteREO Discovery.V8; Carl Zeiss, Oberkochen, Germany), and the tissues were snap-frozen in liquid nitrogen. Total RNA was extracted from each tissue using Sepasol-RNA I (Nacalai Tesque, Kyoto, Japan). The extracted RNA was further purified and treated with TURBO DNase as previously described.^43,44^ RNA quality was assessed using a bioanalyzer (Agilent 2100; Agilent, Santa Clara, CA, USA), confirming that the RNA integrity number (RIN) was 10. RNA sequencing was conducted by Novogene Co., Ltd as contract research *via* NIPPON Genetics Co, Ltd, and raw reads from each tissue were obtained. The resulting reads were aligned to *C*.*robusta* genes (HT version, KY21 gene models) using the Hisat2 algorithm. Gene models were downloaded as “HT.KY21Gene.2.fasta” from the Ghost Database (http://ghost.zool.kyoto-u.ac.jp/default_ht.html). Gene expression levels were calculated as transcripts per million (TPM). Genes that were expressed at levels more than tenfold higher in the OPO compared to other tissues were identified, and their expression levels were further analyzed by Real-time PCR.

### Real-time PCR analysis

Total RNA was extracted from the cerebral ganglion, OPO, spermiduct, oviduct, and ovary as described above. A 500-ng aliquot of DNase-treated total RNA was used for first-strand cDNA synthesis. Real-time PCR was performed using a CFX96 Touch Real-Time PCR Detection System (Bio-Rad Laboratories, Inc., Hercules, CA) as previously described^45^ to validate the RNA-seq data. The primers used for real-time PCR analyses are listed in Table 1. Gene expression levels were normalized to the expression of the ubiquitin-associated domain containing 1 gene (*CiUbac1*, KY21.Chr12.26), which was constitutively expressed among tissues according to the RNA-seq analysis.

**Table 1.**
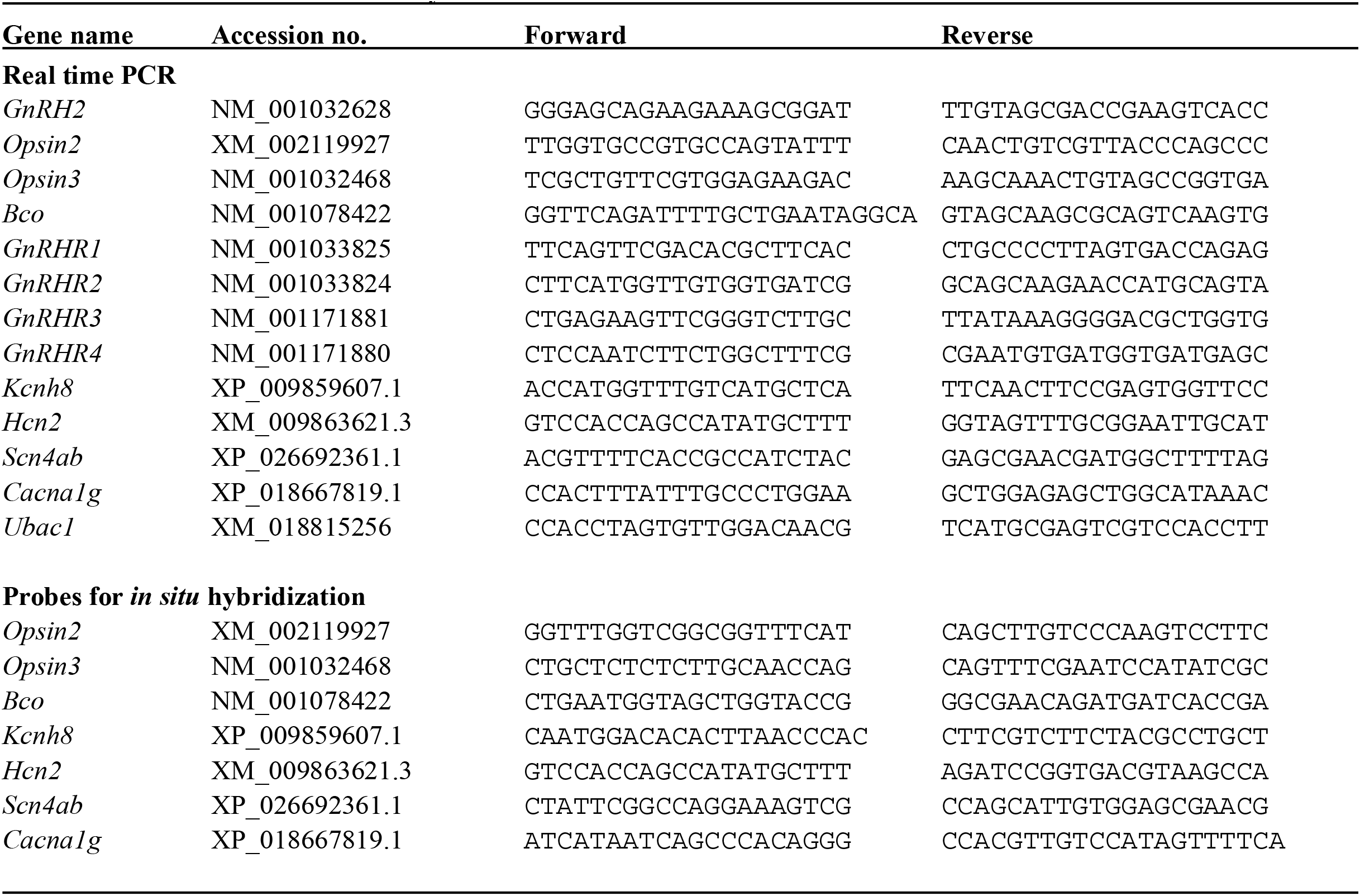
Primer sets used in this study.

### *In situ* hybridization

The RNA probes for *Opsin2* (XM_002119927), *Opsin3* (NM_001032468), and *Bco* (NM_001078422) were designed to span exon-intron junctions. Approximately 300-bp fragments of these genes were inserted into the pCR4-TOPO vector (Thermo Fisher Scientific Inc., Waltham, MA, USA). The plasmids were linearized and used for in vitro transcription with a digoxigenin-labeled RNA labeling kit (Roche Applied Science, Penzberg, Germany). OPO tissues were dissected and fixed in 4% paraformaldehyde in 0.1 M phosphate buffer at 4 °C overnight. The fixed tissues were then soaked in a refrigerated sucrose solution (30% sucrose in PBS) until they sank. They were embedded in Super Cryoembedding Medium-L1 (Leica Microsystems Japan, Tokyo, Japan) and sectioned at 10 µm thickness using a CryoStar NX70 cryostat (Thermo Fisher Scientific Inc., Waltham, MA, USA) at ™20 °C. The sections were mounted onto MAS-coated slides (Matsunami Glass Ind., Ltd., Osaka, Japan). Hybridization, washing, and detection were performed as previously described.^45^ No positive signals were observed when sense probes were used, confirming the specificity of the hybridization. The primer sets used for probe preparation are listed in Table 1. In tissue sections of the OPO, orange-pigmented cells were observed prior to *in situ* hybridization. However, the pigmentation disappeared following the hybridization procedure. To accurately determine the localization of *in situ* hybridization signals relative to the original pigmentation, we captured microscopic images of the OPO both before and after *in situ* hybridization and overlaid the images using Adobe Photoshop Elements 13 (Adobe Systems, San Jose, CA, USA).

## Results

### Observation of spawning behavior

Since continuous light is known to inhibit spawning in *C. robusta*, we hypothesize that dark conditions would stimulate spawning behavior. To test this hypothesis, we placed *C. robusta* under dark conditions and monitored spawning behavior. Spawning began approximately 30 minutes after the onset of darkness (Movie 1). Another video recording revealed that *C. robusta* initiates spawning by releasing sperm, followed by egg release (Movie 2). To examine this sequence more closely, we conducted further observations under a stereomicroscope, focusing on the OPO. These observations confirmed that sperm release precedes egg release (Movie 3). Notably, at the onset of spawning, a knob-like structure on the spermiduct was observed to suppress the oviduct opening. As sperm release progressed, the size of this knob gradually decreased, allowing eggs to begin moving toward release (Movie 3).

### Whole mount observations of transgenic animals

We examined the distribution of *Gnrh1-* and *Gnrh2-*promoter-driven Kaede-positive cells throughout the *C. robusta* body. Figure 1 provides an overview of the distribution of Kaede-positive cells and nerve fibers in the cerebral ganglion, dorsal strand plexus (DSP), OPO, anus, and ovary. In *Gnrh1>Kaede* transgenic animals, Kaede-positive cells were observed in the cerebral ganglion and DSP, with both cells and nerve fibers extending toward the ovary (Fig. 1A-D). In contrast, no projections were observed toward the OPO or anus. These findings are consistent with previous immunohistochemical studies of GnRH peptides^26,36^. In *Gnrh2>Kaede* transgenic animals, Kaede-positive cells were not detected in the cerebral ganglion, but were present in the OPO and anus (Fig. 1 E-G). Nerve fibers appear to project from the OPO to the cerebral ganglion (Fig. 1 A). In *Gnrh2>Kaede* transgenic animals, nerve fibers were not observed projecting to the ovary. Weak nerve projections were detected in the oral siphon in both *Gnrh1>Kaede* and *Gnrh2>Kaede* animals, whereas nerve fibers projected to the atrial siphon only in *Gnrh2>Kaede* animals (Supplemental Fig. 1). Whole-mount observations revealed distinct patterns of cellular and fiber distribution between *Gnrh1* and *Gnrh2* transgenic animals. Interestingly, while *Gnrh2* expression in adult *C. robusta* has been considered to be very low^46^, the present observations suggest specific expression of *Gnrh2* in the OPO and anus, which have not been examined in previous studies.

**Figure 1.**
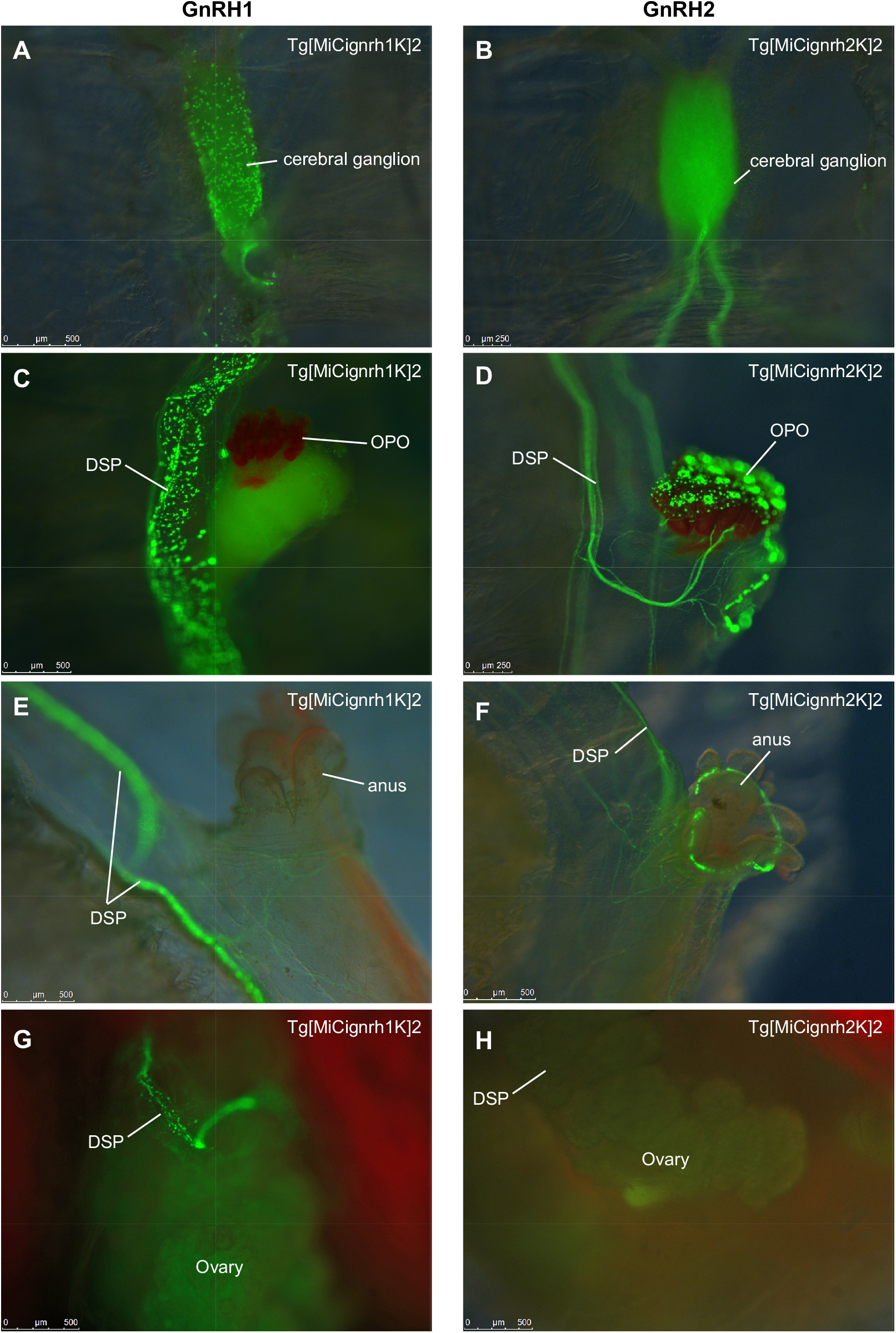
Whole-mount observations of transgenic *C. robusta* expressing Kaede under the control of either the *Gnrh1* or *Gnrh2* promoter. Images were taken using a fluorescence stereo microscope. The names of transgenic lines are indicated at the top-right corner of each panel. (A, B) Cerebral ganglion of *Gnrh1* and *Gnrh2* transgenic lines. (C, D) Dorsal strand plexus (DSP) and orange-pigmented organ (OPO) of *Gnrh1* and *Gnrh2* transgenic lines. (E, F) Anus of *Gnrh1* and *Gnrh2* transgenic lines. (G, H) Ovary and DSP of *Gnrh1* and *Gnrh2* transgenic lines. Scale bars: 500 µm in A, C, E, F, G, and H; 250 µm in B and D.

### Histological analysis of tissue sections of the OPO

Hematoxylin-eosin staining revealed the basic structures of the OPO, spermiduct, and oviduct. The OPO is composed of several papillae, which serve as outlet ports for sperm. These papillae are filled with orange-pigmented cells and are entirely covered by an epithelium layer as reported previously^47^ (Fig. 2). The spermiduct terminates at the OPO, and sperm accumulate in a structure referred to as the spermiduct knob-like structure in this study. The oviduct lies adjacent to the spermiduct, and its opening seems to be pressed down by the spermiduct knob-like structure (Fig. 2). Orange-pigmented cells were observed beneath the epithelial cells, in a region where hemocytes circulate. As shown in Figure 1F, Kaede-positive cells were observed in the OPO of *Gnrh2>Kaede* animals. To further investigate their localization, we examined tissue sections for the distribution of Kaede-positive cells. Fluorescence microscopy revealed that Kaede-positive cells are located within the epithelial layer surrounding the outlet ports of the OPO (Fig. 3). Confocal microscopy further demonstrated that these Kaede-positive cells are neurons, with observable nerve fiber projections (Fig. 3D).

**Figure 2.**
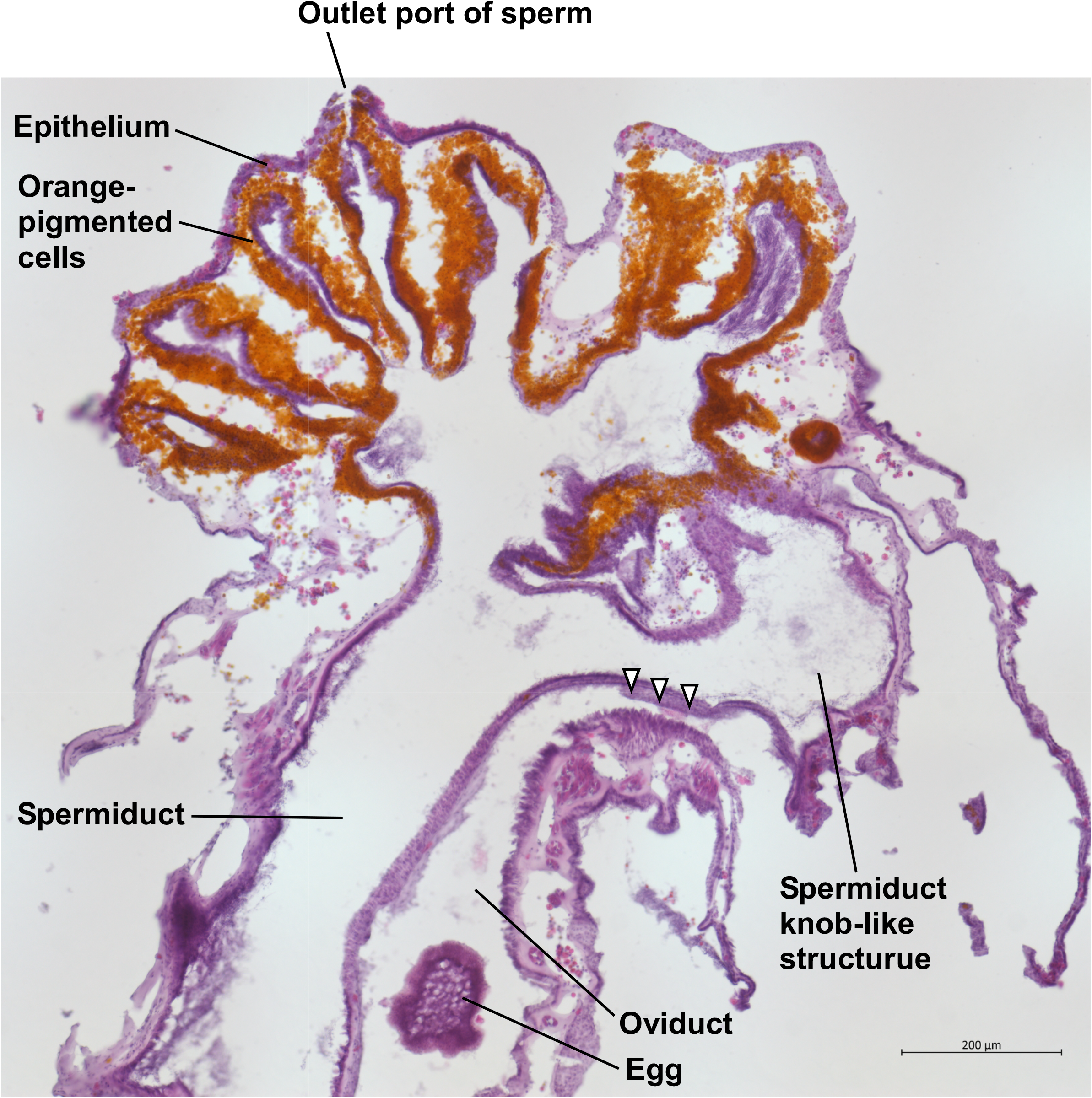
HE staining of the OPO tissue section. The spermiduct terminates at the OPO, which serves as the outlet port for sperm. The OPO consists of multiple papillae filled with orange-pigmented cells. The oviduct terminates beneath the knob-like structure of the spermiduct, and its openings are compressed by this structure (arrowheads). Scale bar: 200 µm.

**Figure 3.**
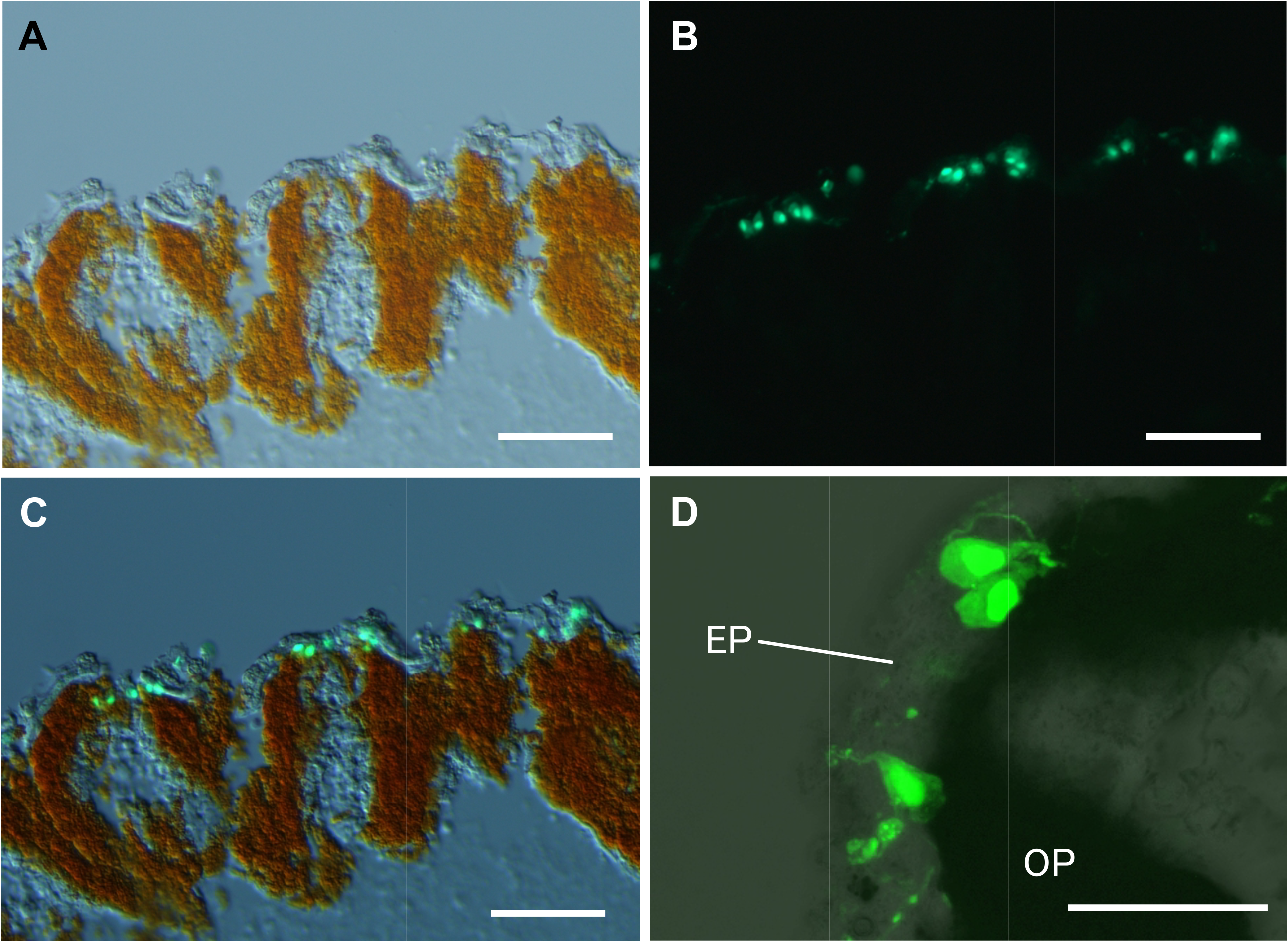
Morphological observations of the OPO section of *Gnrh2* transgenic line (Tg[MiCignrh2K]2). (A) Bright-field image of papillae in the OPO. (B) Dark-field image of the OPO showing *Gnrh2*-positive cells. (C) Merged image of the bright-field and dark-field images. *Gnrh2*-positive cells are observed around the outlet ports of the OPO. (D) Confocal microscopic image showing *Gnrh2*-positive cells located in the epithelium (EP), adjacent to the orange-pigmented cells (OP). Nerve fibers are also visible. Scale bars: 50 µm in A, B, and C; 20 µm in D.

### RNA sequencing and real-time PCR

To investigate gene expression in the OPO, we performed RNA sequencing (RNA-seq) on four tissues: OPO, spermiduct, oviduct, and ovary. RNA-seq yielded approximately 40 to 50 million paired-end reads per tissue. For each dataset, approximately 50,000 transcripts were successfully mapped. The resulting FASTQ files were deposited in the Sequence Read Archive (SRA) under the following accession number: PRJNA1279780. Genes expressed at levels more than tenfold higher in the OPO than in the other tissues were identified and listed. Notably, *Gnrh2* was among the top 100 OPO-specific genes (Supplemental Table 1), consistent with our histological findings based on *Gnrh2* promoter-driven Kaede fluorescence in the OPO of transgenic animals. Interestingly, several photoreception-related genes including *Opsin2, Opsin3*, and beta-carotene-15,15’-monooxygenase (*Bco*), were specifically expressed in the OPO (Supplemental Table 1). In addition, multiple ion channel-related genes were expressed abundantly in the OPO, including potassium/sodium hyperpolarization-activated cyclic nucleotide-gated channel 2 (*Hcn2*)-like, sodium channel protein type 4 subunit alpha B (*Scn4b*)-like, potassium voltage-gated channel subfamily H member 8 (*Kcnh8*)-like, and voltage-dependent T-type calcium channel subunit alpha-1G (*Cacna1*) (Supplemental Table 1). We further examined the expression of these genes by quantitative real-time PCR (qRT-PCR) in the cerebral ganglion, OPO, spermiduct, oviduct, and ovary and found that their expression was almost exclusive in the OPO (Fig. 4). We also analyzed the expression of GnRH receptor genes, including *Gnrhr1* through *Gnrhr4*. In contrast to the OPO-specific genes, the GnRH receptor genes showed relatively high expression in the cerebral ganglion and oviduct, rather than in the OPO (Fig. 4).

**Figurer 4.**
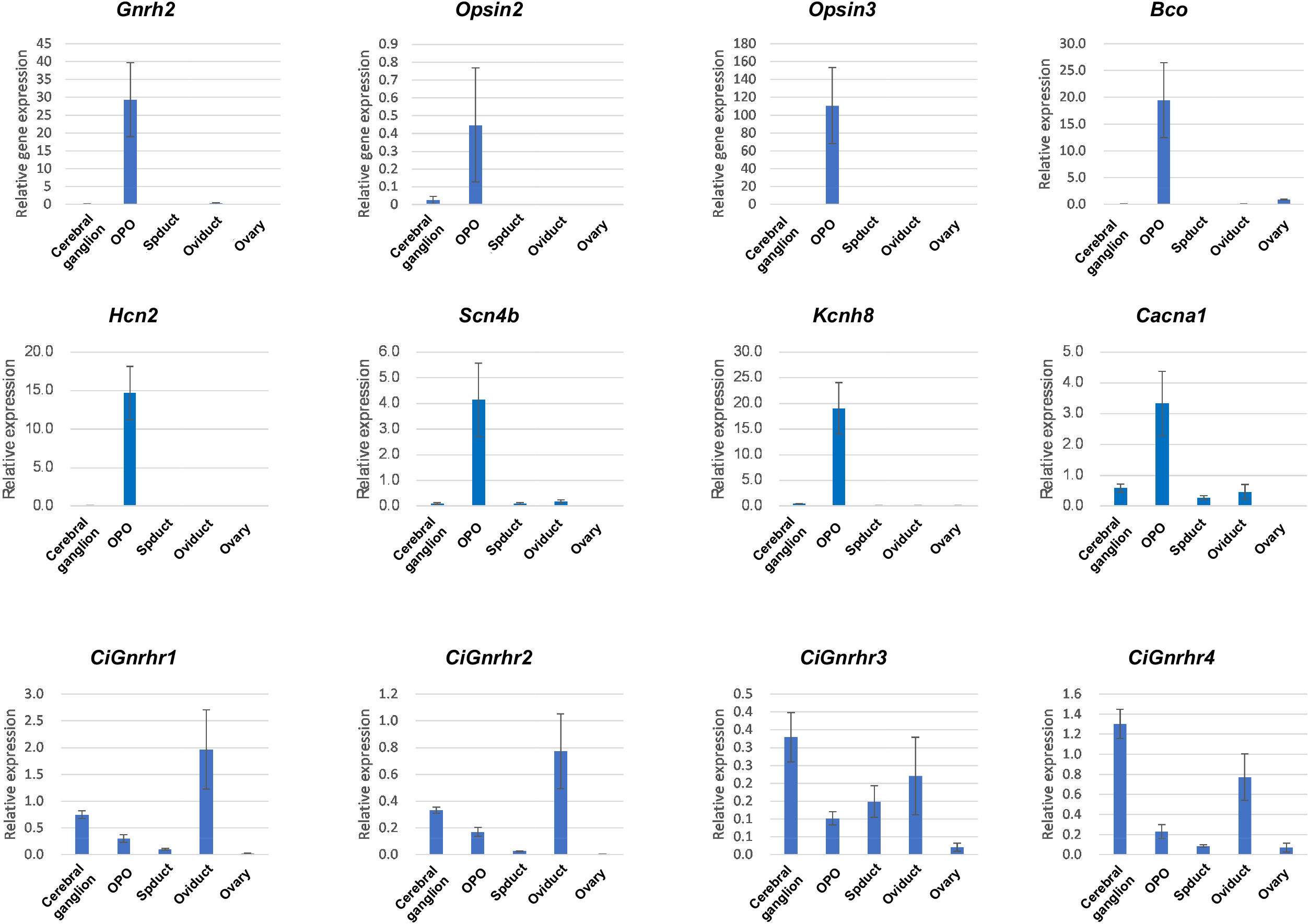
Real-time PCR analysis in the cerebral ganglion, OPO, spermiduct, oviduct, and ovary. Relative expression levels of target genes are shown for each tissue. Data are presented as means ± SEM (*n* = 3).

### *In situ* hybridization

To examine the localization of OPO-specific genes, we performed *in situ* hybridization on OPO tissue sections. As shown in Figure 2, orange-pigmented cells were observed beneath the epithelial layer. *In situ* hybridization signals were detected in a restricted region of the epithelial cells, where those cells are adjacent to the sperm in the papilla of the OPO (Fig. 5). No signal was observed at the distal tip of the OPO opening. Interestingly, all examined genes, including photoreception-related genes and ion channels examined were localized to the same epithelial region (Fig. 5). No hybridization signal was detected using the sense probe, confirming the specificity of the observed signals. Some blue staining was observed in the outer region of the epithelium, which was considered nonspecific (Fig. 5).

**Figure 5.**
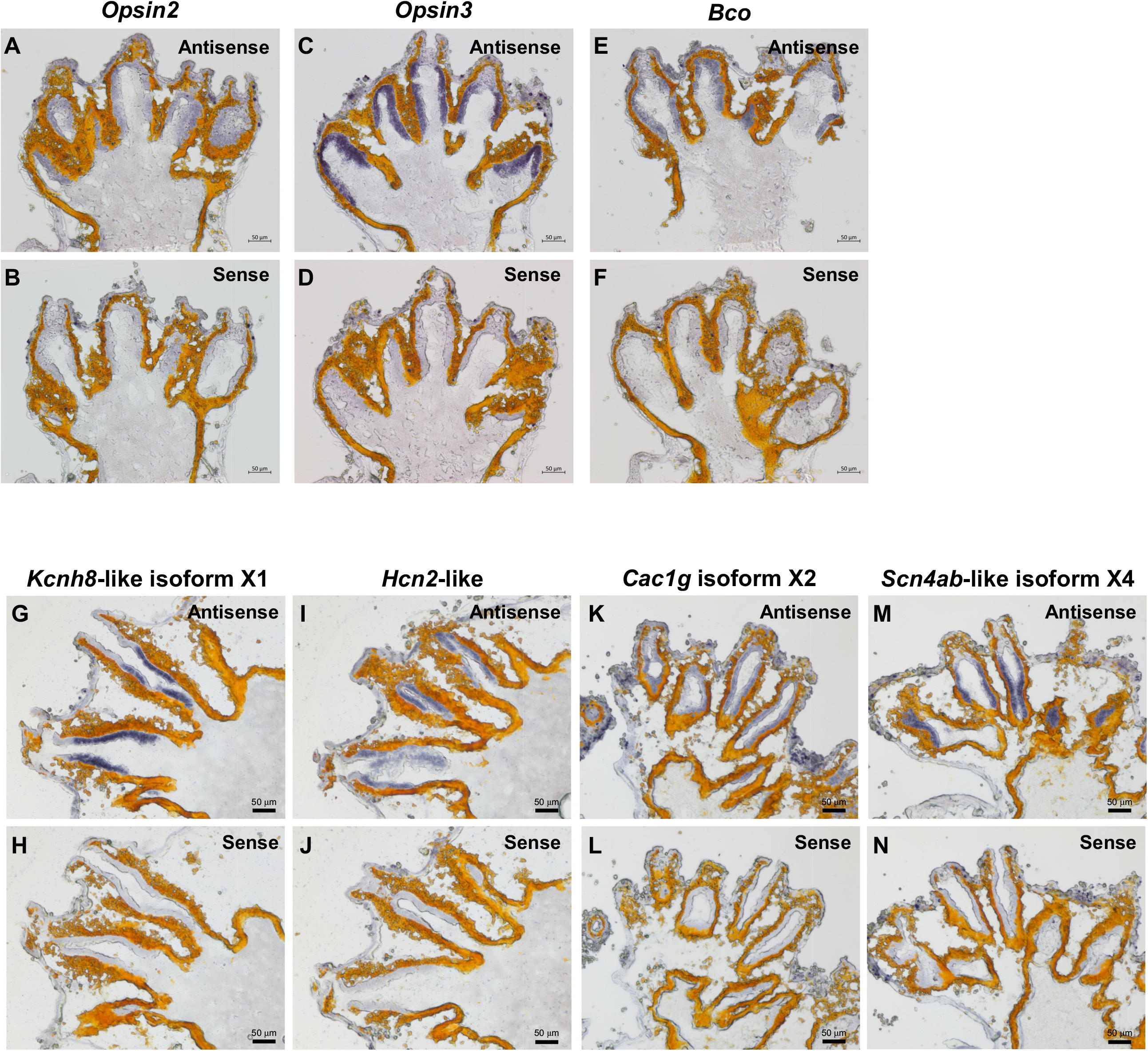
*In situ* hybridization of OPO-specific genes. The upper panels show antisense probe signals for each gene, and lower panels show the corresponding sense controls. mRNA expression is indicated by purple staining. (A, B) *Opsin2*; (C, D) *Opsin3*; (E, F) *Bco*; (G, H) *Kcnh8*-like isoform X1; (I, J) *Hcn2*-like; (K, L) *Cac1g* isoform X2; (M, N) *Scn4ab*-like isoform X4. Scale bars: 50 µm.

## Discussion

In ascidians, *C. intestinalis* and *C. robusta*, which were previously considered to be the same species, have both been used to study spawning behavior. Interestingly, several studies have reported differing light/dark conditions that stimulate spawning in *Ciona*. Some have shown that spawning generally occurs around sunrise.^6-8^ Conversely, other studies have reported spawning during the evening and night-time hours.^9^ Furthermore, one report suggested *Ciona* may spawn and settle at any time of the day.^10^ In our observations, *C. robusta* began spawning approximately 30 minutes after the onset of darkness under laboratory conditions. These results support the view that spawning occurs in *Ciona* occurs during night-time hours.^*9*^ Given that many earlier studies did not distinguish between *C. intestinalis* and *C. robusta*, this lack of taxonomic resolution may have contributed to the discrepancies in previous findings. In light of our results, it may be necessary to re-evaluate the relationship between light–dark stimuli and spawning behavior, particularly in *C. intestinalis*. Until now, the sequence of sperm and egg release during spawning in *C. robusta* has not been thoroughly investigated. In this study, we demonstrated that egg release is physically delayed due to the anatomical arrangement of the spermiduct and oviduct openings, resulting in the release of sperm occurring first. These findings suggest that sperm release in *C. robusta* is actively regulated. In contrast, egg release appears to be passively controlled, suggesting that *Ciona* employs a dexterous strategy which can control the timing of sperm and egg release strictly with the minimum biological cost of active regulation.

In *Ciona*, two gonadotropin-releasing hormone homolog genes, *Gnrh1* and *Gnrh2* have been identified.^26^ Both genes are expressed during embryonic development and in the larval stage.^26,46^ In adult animals, *Gnrh1* is expressed in the neural complex, including the cerebral ganglion, whereas *Gnrh2* is not.^46^ Immunohistochemical studies revealed that GnRH-positive cells are detected in the cerebral ganglion and they project fibers, along with the DSP, to the ovary,^26,36^ suggesting that the GnRH system plays a regulatory role in ovarian functions. Our whole-mount observations of transgenic animals further revealed that the GnRH1-expressing cells are localized in the cerebral ganglion and project nerve fibers to the ovary, supporting the idea that the GnRH1 nervous system may regulate ovarian functions. Interestingly, our observations also showed that GnRH2-expressing cells are localized in the OPO. *Gnrh2*-positive neurons are situated near the outlet ports of the OPO and extend projections to the cerebral ganglion, suggesting that GnRH2 nervous system may be involved in transmitting sperm-release signals to the cerebral ganglion, or in suppressing sperm release under light conditions. To further investigate the role of *Gnrh2* in spawning behavior, we are currently conducting gene knockout experiments.

The present study found key molecules associated with spawning behavior and light/dark conditions. Two opsin genes, *Opsin2* and *Opsin3*, were specifically expressed in the OPO. In a previous phylogenetic analysis, *Opsin2* was grouped within the clade of vertebrate pigments and related opsins, while *Opsin3* was clustered with the human retinal G protein-coupled receptor (RGR), a non-visual opsin that can bind and photoisomerize all-trans-retinal to 11-cis-retinal.^48^ In addition, *Bco* was also expressed in the OPO. All these photoreception-related genes were localized in the innermost region of the OPO, where they are situated adjacent to the orange-pigmented cells. Since *BCO* and *RGR* are expressed in the reginal pigment epithelium (RPE) of mammalian eyes,^49,50^ the innermost region of the OPO may be similar to the RPE in mammals, although it does not form a single-cell layer as in the mammalian RPE.^51^ It is particularly interesting that *Opsin2*, a photoreceptor gene, was expressed in the same region as *Bco* and *Opsin3*, given that in mammalian eyes, RPE cells are functionally or physically separated from photoreceptor cells.^50^ Although *Ciona* is positioned as the sister group of vertebrates and its organs are less differentiated, these findings suggests that the OPO functions as a photoreceptive organ involved in spawning behavior in response to light/dark cues. In *Ciona* larvae, *Opsin1* a paralog of *Opsin2*, is expressed in the outer segment of photoreceptor cells, whereas *Opsin2* is not found in the ocellus photoreceptor cells but instead is expressed in the brain vesicle and oral siphon rudiment.^48^ *Opsin3* is localized in both the ocellus photoreceptor cells and surrounding non-photoreceptor cells in the brain vesicle.^48,52^ These photoreceptive systems in *Ciona* larvae are important for the regulation of swimming behavior,^53,54^ and likely contribute to the regulation of spawning behavior in adult *Ciona*, although the developmental transition of this system remains to be elucidated.

In this study, we also focused on the ion channels in the RNA-seq data, as they are known to play important roles in visual function in vertebrates.^55,56^ Several putative ion channel genes, including *Hcn2*-like, *Scn4b*-like, *Kcnh8*-like, and *Cacna1*, were specifically expressed in the same region of the photoreceptive site in the OPO. Although we have not yet conducted double-staining analyses, photoreceptor-related genes and ion channel genes appear to be co-expressed in the same cells. In mammals, phototransduction pathway involved in non-image-forming vision depends on HCN channels.^56^ In retinoblastoma cells, *CACNA1G* is expressed but is downregulated after differentiation.^57^ These findings suggest that *Hcn2*-like and *Cacna1* may be involved in phototransduction in the OPO in a manner similar to their roles in mammals. In contrast, to the best of our knowledge, there is no evidence that *Scn4b* and *Kcnh8* participate in vertebrate photoreceptor systems. These two ion channels may therefore have *Ciona*-specific roles within the photoreceptive system in the OPO.

In summary, this study revealed that spawning behavior in *C. robusta* is regulated by light/dark conditions, with spawning initiated after the onset of darkness. Morphological, gene expression, and histological analyses indicate that the OPO functions as a photoreceptive organ, and that the GnRH2, rather than GnRH1, neural system participates in the regulation of spawning behavior. Further investigations will be necessary to elucidate potential interactions among GnRH2, photoreception-related genes, and ion channels in the context of gamete release regulation in the OPO. Functional analyses of the identified genes and neural circuits are currently in progress and will be essential for fully understanding the mechanism of light-mediated spawning in ascidians.

## Acknowledgments

We gratefully acknowledge Prof. Fumihiko Sato for his fruitful comments on the manuscript. We thank the National Bio-Resource Project (NBRP) for providing ascidians. This study was supported by JSPS Grants-in-Aid for Scientific Research (KAKENHI); Grant Numbers 22K06327 and 25K09712 to T.O., and 23K27185 to T.G.K.

## Author contributions

T.O. designed the study; T.O., S.M., A.S., A.W., Y.M., I.S.S., Y.S., T.G.K., and H.S. performed the research; T.O., S.M., A.S., Y.S., T.G.K., and H.S. wrote, reviewed, and edited the paper. Y.S. produced and provided transgenic animals *via* NBRP.

## Competing interests

The authors declare no competing interests.

**Correspondence and requests** for materials should be addressed to T.O.

---

**Supplemental Figure 1. Whole-mount observations of transgenic *C. robusta* expressing Kaede under the control of either the *Gnrh1* or *Gnrh2* promoter**. Images were taken using a fluorescence stereo microscope. The names of transgenic lines are indicated at the top-right corner of each panel. The right panels show dark-field images, and the left panels show merged images of bright-field and dark-field views. (Aa, Ab) Oral siphon of *Gnrh1* transgenic line. (Ac, Ad) Oral siphon of *Gnrh2* transgenic line. (Be, Bf) Atrial siphon of *Gnrh1* transgenic line. (Bg, Bh) Atrial siphon of *Gnrh2* transgenic line. Scale bars: 1 mm.

**Movies**

Movie 1, 2, and 3 are available in the Zenodo repository under the following doi number: 10.5281/zenodo.15734177.

## Notes

https://doi.org/10.5281/zenodo.15734176

### Competing Interest Statement

The authors have declared no competing interest.

### Summary of Updates

Contact information of an author is updated.

