## Supplementary figures and images for "Gamete release in *Ciona robusta*: roles of gonadotropin-releasing hormone and the photoreception system"

### Supplemental Fig 1.pdf

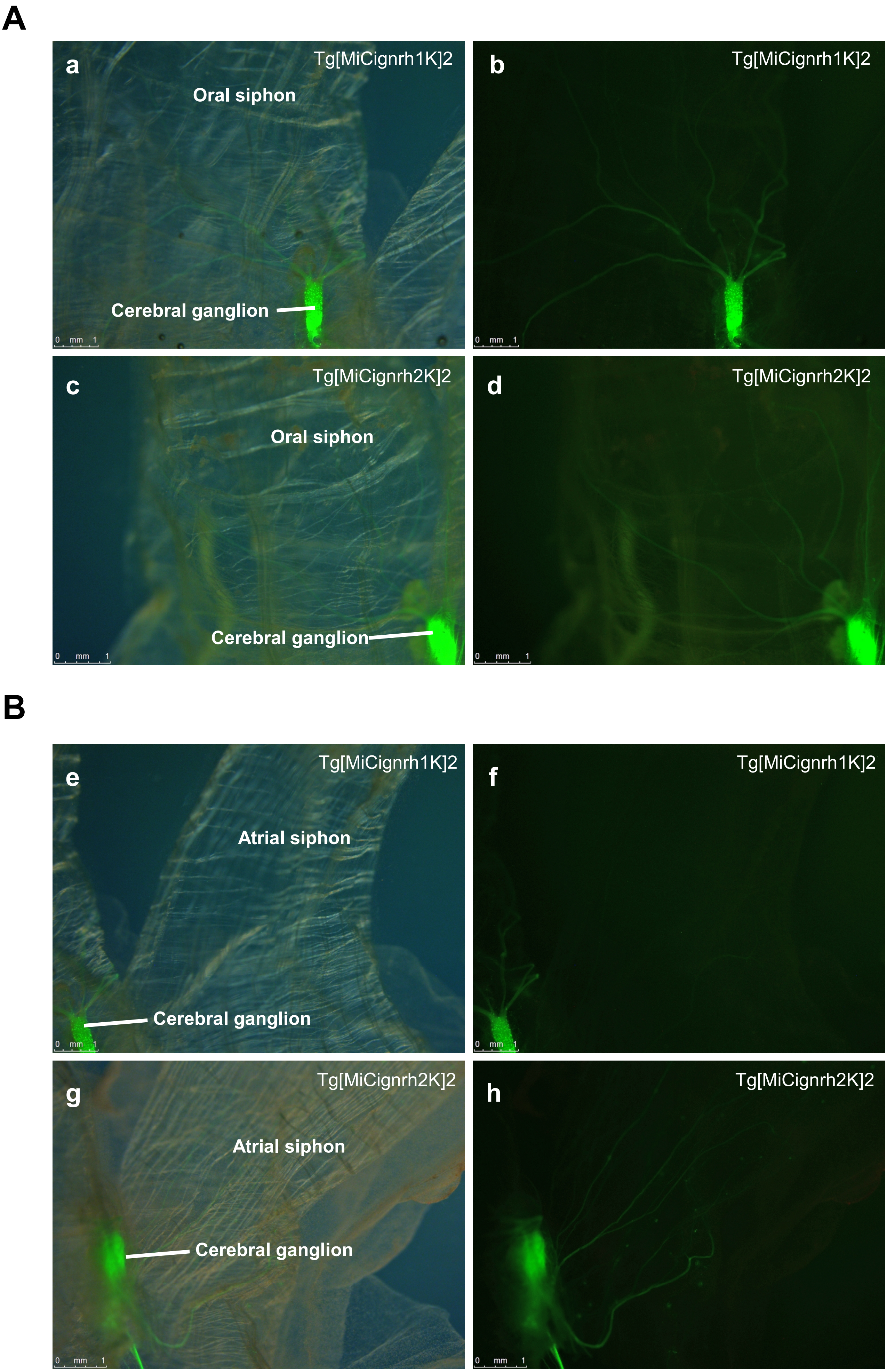
