## Supplemental Table 1 for "Gamete release in *Ciona robusta*: roles of gonadotropin-releasing hormone and the photoreception system": Supplemental Table 1.pdf

**Supplemental Table 1. Top 100 of OPO specific genes**

| gene_id | transcript_id(s) | length | OPO_TPM | Spduct_TPM | Oviduct_TPM | Ovary_TPM | OPO vs Spduct | OPO vs Oviduct | OPO vs Ovary | Annotation |
|---|---|---|---|---|---|---|---|---|---|---|
| KY21.Chr1.1819 | KY21:KY21.Chr1.1819 | 973.96 | 45791.2 | 9.5 | 2.43 | 0.44 | 4820.13 | 18844.12 | 104070.93 | uncharacterized protein (XP_026692539.1) |
| KY21.Chr12.157 | KY21:KY21.Chr12.157 | 1321 | 6948.44 | 0.89 | 0.31 | 6.37 | 7807.24 | 22414.32 | 1090.81 | Opsin3 (retinal G-protein-coupled receptor) (NP_001027640.1) |
| KY21.Chr9.932 | KY21:KY21.Chr9.932.v | 2046.12 | 2752.28 | 0.09 | 0 | 51.45 | 30580.89 | N.A. | 53.49 | beta-carotene-15,15'-monooxygenase (NP_001071890.1) |
| KY21.Chr1.1938 | KY21:KY21.Chr1.1938 | 2114 | 1859.27 | 55.99 | 18.09 | 47.24 | 33.21 | 102.78 | 39.36 | RIC-8(Synembryn) (XP_002121749.1) |
| KY21.Chr6.80 | KY21:KY21.Chr6.80.v1 | 2508 | 1742.76 | 0.21 | 87.78 | 1.97 | 8298.86 | 19.85 | 884.65 | Ci-meta2 (NP_001027777.1) |
| KY21.Chr9.123 | KY21:KY21.Chr9.123.v | 938 | 1664.74 | 0 | 7.74 | 0 | N.A. | 215.08 | N.A. | GnRH-2 (NP_001027800.2) |
| KY21.Chr14.721 | KY21:KY21.Chr14.721 | 2813 | 1342.74 | 0.03 | 51.24 | 3.82 | 44758.00 | 26.20 | 351.50 | Uncharacterized protein (unknown) |
| KY21.Chr9.729 | KY21:KY21.Chr9.729.v | 1547.59 | 1274.89 | 110.31 | 104.19 | 95.86 | 11.56 | 12.24 | 13.30 | ubiquitin-conjugating enzyme E2 E1 (XP_002129445.1) |
| KY21.Chr4.922 | KY21:KY21.Chr4.922.v | 2216 | 835.04 | 1.26 | 0 | 0.09 | 662.73 | N.A. | 9278.22 | potassium/sodium hyperpolarization-activated cyclic nucleotide-gated channel 2-like (XP_009861923.1) |
| KY21.Chr11.401 | KY21:KY21.Chr11.401 | 1923 | 822.38 | 0.57 | 25.12 | 28.9 | 1442.77 | 32.74 | 28.46 | coadhesin isoform X1 (XP_002130909.1) |
| KY21.Chr13.83 | KY21:KY21.Chr13.83.v | 3093.12 | 784.97 | 0.48 | 46.75 | 0.48 | 1635.35 | 16.79 | 1635.35 | uncharacterized (XP_002130436.1) |
| KY21.Chr6.486 | KY21:KY21.Chr6.486.v | 1051.9 | 724.42 | 23.1 | 26.25 | 4.92 | 31.36 | 27.60 | 147.24 | tumor protein p53-inducible protein 11 (XP_002130208.1) |
| KY21.Chr1.2012 | KY21:KY21.Chr1.2012 | 2623.48 | 705.03 | 0.13 | 17.17 | 0.48 | 5423.31 | 41.06 | 1468.81 | Epi-1 precursor (NP_001027612.1) |
| KY21.Chr14.34 | KY21:KY21.Chr14.34.v | 1049 | 654.9 | 0.4 | 0 | 0 | 1637.25 | N.A. | N.A. | Opsin2 (pinopsin-like) (XP_002119963.1) |
| KY21.Chr3.1599 | KY21:KY21.Chr3.1599 | 234 | 567.99 | 15.76 | 0 | 0 | 36.04 | N.A. | N.A. | sodium channel protein type 4 subunit alpha B-like isoform X4 (XP_026692361.1) |
| KY21.Chr14.535 | KY21:KY21.Chr14.535 | 2137.18 | 565.81 | 0.13 | 6.33 | 0 | 4352.38 | 89.39 | N.A. | deleted in malignant brain tumors 1 protein (XP_002131041.1) |
| KY21.Chr9.50 | KY21:KY21.Chr9.50.v1 | 3822.42 | 505.19 | 3.78 | 2.5 | 2.35 | 133.65 | 202.08 | 214.97 | potassium voltage-gated channel subfamily H member 8-like isoform X1 (XP_009859607.1) |
| KY21.Chr5.737 | KY21:KY21.Chr5.737.v | 1659 | 494.3 | 18.4 | 16.54 | 27.13 | 26.86 | 29.89 | 18.22 | ras-related protein Rab-27B (XP_002127854.1) |
| KY21.Chr3.1271 | KY21:KY21.Chr3.1271 | 1838.07 | 464.35 | 4.26 | 4.55 | 3.76 | 109.00 | 102.05 | 123.50 | intermediate filament protein IF-C (NP_001027953.2) |
| KY21.Chr2.441 | KY21:KY21.Chr2.441.v | 1740.22 | 437.8 | 12.83 | 17.48 | 1.91 | 34.12 | 25.05 | 229.21 | uncharacterized (XP_026696328.1) |
| KY21.Chr12.240 | KY21:KY21.Chr12.240 | 666.12 | 418.13 | 0.41 | 4.22 | 4.31 | 1019.83 | 99.08 | 97.01 | lymphocyte antigen 6D-like isoform X1 (XP_002122388.1) |
| KY21.Chr3.1600 | KY21:KY21.Chr3.1600 | 6214 | 416.82 | 13.9 | 14.69 | 4.42 | 29.99 | 28.37 | 94.30 | sodium channel protein type 4 subunit alpha B-like isoform X4 (XP_026692361.1) |
| KY21.Chr2.817 | KY21:KY21.Chr2.817.v | 2467.79 | 413.87 | 0.04 | 14.06 | 0 | 10346.75 | 29.44 | N.A. | cubilin isoform X1 (XP_018673033.1) |
| KY21.Chr8.1283 | KY21:KY21.Chr8.1283 | 1706.61 | 403.35 | 18.92 | 16.23 | 29.69 | 21.32 | 24.85 | 13.59 | cytochrome b561 (NP_001041449.1) |
| KY21.Chr1.1206 | KY21:KY21.Chr1.1206 | 1858 | 327.27 | 0.94 | 4.41 | 0.23 | 348.16 | 74.21 | 1422.91 | CD94/NKR-P1-like protein precursor (NP_001027956.2) |
| KY21.Chr1.1671 | KY21:KY21.Chr1.1671 | 1550 | 298.17 | 0.12 | 0 | 0 | 2484.75 | N.A. | N.A. | transcription factor protein isoform X1 (XP_026694905.1) |
| KY21.Chr9.276 | KY21:KY21.Chr9.276.v | 305 | 289.37 | 0 | 0 | 0 | N.A. | N.A. | N.A. | uncharacterized (XR_717376.3) |
| KY21.Chr13.359 | KY21:KY21.Chr13.359 | 488 | 288.64 | 0.64 | 1.17 | 0 | 451.00 | 246.70 | N.A. | uncharacterized (XR_003396510.1) |
| KY21.Chr1.1818 | KY21:KY21.Chr1.1818 | 319.57 | 278.68 | 0 | 0.6 | 0 | N.A. | 464.47 | N.A. | uncharacterized (XR_182027.4) |
| KY21.Chr7.977 | KY21:KY21.Chr7.977.v | 2627.12 | 270.97 | 5.1 | 15.38 | 5.29 | 53.13 | 17.62 | 51.22 | sushi, von Willebrand factor type A, EGF and pentraxin domain-containing protein 1 isoform X1 (XP_002129375.1) |
| KY21.Chr8.986 | KY21:KY21.Chr8.986.v | 1014.03 | 268.11 | 16.2 | 3.99 | 2.62 | 16.55 | 67.20 | 102.33 | bromodomain-containing protein 8 (XM_009861063.2) |
| KY21.Chr1.1498 | KY21:KY21.Chr1.1498 | 1154.43 | 263.26 | 18.86 | 12.63 | 9.87 | 13.96 | 20.84 | 26.67 | transmembrane emp24 domain-containing protein eca-like (XM_002131471.4) |
| KY21.Chr9.802 | KY21:KY21.Chr9.802.v | 1990 | 234.95 | 0.05 | 1.9 | 0.42 | 4699.00 | 123.66 | 559.40 | glial fibrillary acidic protein (NM_001032544.2) |
| KY21.Chr12.1050 | KY21:KY21.Chr12.105 | 1936 | 225.78 | 1.11 | 0.54 | 0.89 | 203.41 | 418.11 | 253.69 | hemocyanin, beta-C chain unit D (XM_002119639.5) |
| KY21.Chr6.74 | KY21:KY21.Chr6.74.v1 | 2667.42 | 223.94 | 0 | 19.3 | 1 | N.A. | 11.60 | 223.94 | uncharacterized (XM_002124392.4) |
| KY21.Chr12.987 | KY21:KY21.Chr12.987 | 3098 | 222.9 | 7.63 | 1.62 | 3.96 | 29.21 | 137.59 | 56.29 | metalloendopeptidase homolog PEX (XM_002128500.5) |
| KY21.Chr14.172 | KY21:KY21.Chr14.172 | 2249.18 | 214.37 | 1.14 | 2.43 | 0.21 | 188.04 | 88.22 | 1020.81 | complement component C6 (XM_002130752.5) |
| KY21.Chr7.131 | KY21:KY21.Chr7.131.v | 1693.23 | 214.1 | 3.76 | 6.8 | 1.57 | 56.94 | 31.49 | 136.37 | kappa opioid receptor-like (NP_001231977.1) |
| KY21.Chr1.2021 | KY21:KY21.Chr1.2021 | 1251 | 209.2 | 10.87 | 0 | 0.63 | 19.25 | N.A. | 332.06 | actin, muscle (XP_002123216.1) |
| KY21.Chr13.361 | KY21:KY21.Chr13.361 | 4402 | 205.45 | 3.4 | 1.21 | 4.49 | 60.43 | 169.79 | 45.76 | uncharacterized (XP_002124752.2) |
| KY21.Chr10.677 | KY21:KY21.Chr10.677 | 980 | 204.16 | 0.22 | 4.04 | 0 | 928.00 | 50.53 | N.A. | uncharacterized (XP_018669232.1) |
| KY21.UAContig53.14 | KY21:KY21.UAContig5 | 216 | 192.5 | 0 | 0 | 8.97 | N.A. | N.A. | 21.46 | 28S ribosomal RNA (AF212177.1) |
| KY21.Chr10.714 | KY21:KY21.Chr10.714 | 475 | 181.67 | 2.52 | 5.14 | 0.39 | 72.09 | 35.34 | 465.82 | uncharacterized (XR_182129.4) |
| KY21.Chr1.151 | KY21:KY21.Chr1.151.v | 1541 | 176.25 | 0.06 | 2.94 | 0 | 2937.50 | 59.95 | N.A. | uncharacterized (XP_026693699.1) |
| KY21.Chr6.485 | KY21:KY21.Chr6.485.v | 727.01 | 165.8 | 15.58 | 9.51 | 0.56 | 10.64 | 17.43 | 296.07 | uncharacterized (XP_002130314.1) |
| KY21.Chr7.745 | KY21:KY21.Chr7.745.v | 2197 | 165.57 | 1.8 | 12.17 | 0 | 91.98 | 13.60 | N.A. | distal-less isoform X1 (XP_018667881.1) |
| KY21.Chr9.67 | KY21:KY21.Chr9.67.v1 | 1292 | 158 | 5.33 | 6.51 | 0.48 | 29.64 | 24.27 | 329.17 | uncharacterized (XP_002126911.1) |
| KY21.Chr2.933 | KY21:KY21.Chr2.933.v | 2566 | 157.71 | 14.29 | 6.4 | 0.16 | 11.04 | 24.64 | 985.69 | transforming growth factor beta superfamily signaling ligand (NP_001071664.1) |
| KY21.Chr4.338 | KY21:KY21.Chr4.338.v | 3281.42 | 154.14 | 4.88 | 11.56 | 0.45 | 31.59 | 13.33 | 342.53 | sushi, von Willebrand factor type A, EGF and pentraxin domain-containing protein 1-like (XP_002128463.2) |
| KY21.Chr1.1488 | KY21:KY21.Chr1.1488 | 1520.82 | 153.81 | 2 | 10.07 | 1.72 | 76.91 | 15.27 | 89.42 | chymotrypsin-like elastase family member 2A (XP_002131650.1) |
| KY21.Chr10.81 | KY21:KY21.Chr10.81.v | 2256.31 | 153.55 | 0.16 | 7.19 | 0 | 959.69 | 21.36 | N.A. | transmembrane protease serine 9 (XP_002122383.1) |
| KY21.Chr1.1534 | KY21:KY21.Chr1.1534 | 1209.5 | 149.08 | 0.08 | 1.98 | 0.05 | 1863.50 | 75.29 | 2981.60 | GDP-L-fucose synthase (XP_002124066.1) |
| KY21.Chr1.1896 | KY21:KY21.Chr1.1896 | 697.49 | 135.87 | 1.49 | 11.09 | 1.25 | 91.19 | 12.25 | 108.70 | uncharacterized (XP_009862496.1) |
| KY21.Chr3.261 | KY21:KY21.Chr3.261.v | 1400 | 134.2 | 5.59 | 0.96 | 0.94 | 24.01 | 139.79 | 142.77 | protein APCDD1-like isoform X1 (XP_002123300.2) |
| KY21.Chr4.187 | KY21:KY21.Chr4.187.v | 1354 | 128.96 | 0 | 0.04 | 0 | N.A. | 3224.00 | N.A. | testisin-like isoform X2 (XP_018667230.1) |
| KY21.Chr3.972 | KY21:KY21.Chr3.972.v | 12107 | 125.05 | 8.86 | 3.97 | 5.47 | 14.11 | 31.50 | 22.86 | transformation/transcription domain-associated protein isoform X1 (XP_018666778.1) |
| KY21.Chr12.536 | KY21:KY21.Chr12.536 | 1570 | 120.15 | 0.06 | 1.56 | 0 | 2002.50 | 77.02 | N.A. | lipopolysaccharide-binding protein-like (XP_002127342.1) |
| KY21.Chr10.784 | KY21:KY21.Chr10.784 | 1310 | 115.7 | 0.23 | 0.09 | 0.26 | 503.04 | 1285.56 | 445.00 | claudin-1-like isoform (XP_026692137.1) |
| KY21.Chr4.340 | KY21:KY21.Chr4.340.v | 5296.07 | 112.54 | 3.17 | 7.95 | 0.7 | 35.50 | 14.16 | 160.77 | sushi, von Willebrand factor type A, EGF and pentraxin domain-containing protein 1 (XP_009858296.1) |
| KY21.Chr1.566 | KY21:KY21.Chr1.566.v | 1596.3 | 110.74 | 0.06 | 0.11 | 0 | 1845.67 | 1006.73 | N.A. | serine/arginine repetitive matrix protein 4 (XP_026694906.1) |
| KY21.Chr6.621 | KY21:KY21.Chr6.621.v | 1235 | 107.63 | 10.38 | 10.18 | 8.53 | 10.37 | 10.57 | 12.62 | uncharacterized (XP_002130877.1) |
| KY21.Chr5.680 | KY21:KY21.Chr5.680.v | 2256.87 | 106.72 | 4.81 | 4.01 | 2.28 | 22.19 | 26.61 | 46.81 | uncharacterized (XP_026690412.1) |
| KY21.Chr2.251 | KY21:KY21.Chr2.251.v | 2021 | 104.66 | 0.45 | 2.46 | 4.94 | 232.58 | 42.54 | 21.19 | collagen alpha-1(XII) chain (XP_009857560.1) |
| KY21.Chr8.330 | KY21:KY21.Chr8.330.v | 3112 | 103.45 | 0.17 | 1.13 | 0.58 | 608.53 | 91.55 | 178.36 | uncharacterized (XP_002119202.1) |
| KY21.Chr3.87 | KY21:KY21.Chr3.87.v1 | 6111 | 102.87 | 5.76 | 1.71 | 0.26 | 17.86 | 60.16 | 395.65 | C3 and PZP-like alpha-2-macroglobulin domain-containing protein 8 (XP_009861615.2) |
| KY21.Chr1.586 | KY21:KY21.Chr1.586.v | 1094.7 | 100.24 | 2.48 | 7.13 | 0.17 | 40.42 | 14.06 | 589.65 | uncharacterized (XP_026689360.1) |
| KY21.Chr6.493 | KY21:KY21.Chr6.493.v | 1134 | 96.34 | 0.54 | 2.16 | 0 | 178.41 | 44.60 | N.A. | uncharacterized (XP_002129609.1) |
| KY21.Chr3.1410 | KY21:KY21.Chr3.1410 | 712 | 93.64 | 0.85 | 2.35 | 0.39 | 110.16 | 39.85 | 240.10 | basic phospholipase A2 nigroxin A-like isoform X3 (XP_026689604.1) |
| KY21.Chr5.438 | KY21:KY21.Chr5.438.v | 550.31 | 93.12 | 5.35 | 4.17 | 0.46 | 17.41 | 22.33 | 202.43 | uncharacterized (XP_002127290.2) |
| KY21.Chr1.423 | KY21:KY21.Chr1.423.v | 1287 | 92.98 | 0.31 | 1.2 | 0.7 | 299.94 | 77.48 | 132.83 | uncharacterized (XP_002127284.1) |
| KY21.Chr6.290 | KY21:KY21.Chr6.290.v | 1929 | 91.35 | 1.66 | 1.31 | 0.08 | 55.03 | 69.73 | 1141.88 | low-density lipoprotein receptor-related protein 2-like (XP_004226169.1) |
| KY21.Chr3.848 | KY21:KY21.Chr3.848.v | 1606 | 87.38 | 2.05 | 7.69 | 0.17 | 42.62 | 11.36 | 514.00 | putative ferric-chelate reductase 1 (XP_018673419.1) |
| KY21.Chr6.429 | KY21:KY21.Chr6.429.v | 7350 | 86.56 | 7.13 | 5.32 | 1.29 | 12.14 | 16.27 | 67.10 | voltage-dependent T-type calcium channel subunit alpha-1G isoform X2 (XP_018667819.1) |
| KY21.Chr11.1240 | KY21:KY21.Chr11.124 | 2135.2 | 81.14 | 4.65 | 5.1 | 1.67 | 17.45 | 15.91 | 48.59 | stabilin-2-like isoform X3 (XP_002123487.1) |
| KY21.Chr1.2025 | KY21:KY21.Chr1.2025 | 1652.84 | 79.65 | 1.5 | 1.77 | 1.94 | 53.10 | 45.00 | 41.06 | alpha-tectorin-like (XP_002120027.1) |
| KY21.Chr3.1434 | KY21:KY21.Chr3.1434 | 1693.62 | 78.97 | 6.16 | 6.29 | 3.38 | 12.82 | 12.55 | 23.36 | oxidoreductase NAD-binding domain-containing protein 1-like (XM_026833808.1) |
| KY21.Chr8.316 | KY21:KY21.Chr8.316.v | 5309.45 | 77.92 | 2.11 | 4.32 | 0.59 | 36.93 | 18.04 | 132.07 | SCO-spondin (LOC100180388), transcript variant X1, mRNA (XM_026835716.1) |
| KY21.Chr10.515 | KY21:KY21.Chr10.515 | 2410 | 77.89 | 0.07 | 0.02 | 0.53 | 1112.71 | 3894.50 | 146.96 | potassium/sodium hyperpolarization-activated cyclic nucleotide-gated channel 3-like isoform X3 (XP_026692281.1) |
| KY21.Chr2.813 | KY21:KY21.Chr2.813.v | 2752 | 76.49 | 4.43 | 0.94 | 0.09 | 17.27 | 81.37 | 849.89 | tolloid-like protein 2 (LOC100178850), transcript variant X1, mRNA (XM_002128704.4) |
| KY21.Chr8.1155 | KY21:KY21.Chr8.1155 | 1157.44 | 75.62 | 0.17 | 2.72 | 0.35 | 444.82 | 27.80 | 216.06 | uncharacterized (XM_002129511.4) |
| KY21.Chr2.1205 | KY21:KY21.Chr2.1205 | 3031 | 75.57 | 5.83 | 4.63 | 0.03 | 12.96 | 16.32 | 2519.00 | plasminogen (LOC100176264), mRNA (XM_002131067.4) |
| KY21.Chr11.1056 | KY21:KY21.Chr11.105 | 2469 | 75.5 | 3.45 | 4.35 | 0.04 | 21.88 | 17.36 | 1887.50 | kinesin-like protein KIF26B (LOC100178540), mRNA (XM_018813908.2) |
| KY21.Chr2.1031 | KY21:KY21.Chr2.1031 | 1296 | 74.73 | 0.08 | 0.14 | 0 | 934.13 | 533.79 | N.A. | msxb homeoprotein (msxb), mRNA (NM_001032496.1) |
| KY21.Chr7.518 | KY21:KY21.Chr7.518.v | 201 | 73.85 | 0 | 0 | 0 | N.A. | N.A. | N.A. | iron-sulfur cluster assembly scaffold protein IscU-like (XM_026835216.1) |
| KY21.Chr14.583 | KY21:KY21.Chr14.583 | 1391.71 | 73.34 | 0.14 | 0 | 4.31 | 523.86 | N.A. | 17.02 | uncharacterized (XP_002128942.1) |
| KY21.Chr11.1124 | KY21:KY21.Chr11.112 | 1171 | 72.93 | 2.75 | 0.98 | 0.98 | 26.52 | 74.42 | 74.42 | uncharacterized (XP_026692554.1) |
| KY21.Chr9.1009 | KY21:KY21.Chr9.1009 | 246 | 71.67 | 0 | 5.88 | 0 | N.A. | 12.19 | N.A. | zinc finger (MYND)-2 (zf(mynd)-2), mRNA (XM_026835846.1) |
| KY21.Chr11.583 | KY21:KY21.Chr11.583 | 1832.85 | 70.51 | 2.08 | 3.52 | 0 | 33.90 | 20.03 | N.A. | P-selectin-like (XP_002125358.1) |
| KY21.Chr1.491 | KY21:KY21.Chr1.491.v | 1422.25 | 70.22 | 0 | 0.98 | 0 | N.A. | 71.65 | N.A. | uncharacterized (XP_002127176.3) |
| KY21.Chr1.494 | KY21:KY21.Chr1.494.v | 2412 | 69.46 | 0 | 0 | 0.15 | N.A. | N.A. | 463.07 | E-selectin (XP_009861089.1) |
| KY21.Chr2.815 | KY21:KY21.Chr2.815.v | 1701.85 | 68.3 | 0.66 | 1.45 | 0.31 | 103.48 | 47.10 | 220.32 | tolloid-like protein 2 (XP_026696666.1) |
| KY21.Chr14.662 | KY21:KY21.Chr14.662 | 1389 | 67.15 | 0 | 3.06 | 0 | N.A. | 21.94 | N.A. | collagen alpha-1(XII) chain-like (LOC100181428), mRNA (XM_002124374.5) |
| KY21.Chr2.89 | KY21:KY21.Chr2.89.v1 | 2497 | 66.96 | 0.93 | 0.26 | 5.29 | 72.00 | 257.54 | 12.66 | synaptotagmin-like protein 1 (LOC104265437), transcript variant X2, mRNA (XM_026840896.1) |
| KY21.Chr3.1611 | KY21:KY21.Chr3.1611 | 2723 | 65.53 | 0.97 | 1.01 | 2.4 | 67.56 | 64.88 | 27.30 | fibronectin (LOC100181231), transcript variant X3, mRNA (XM_026840138.1) |
| KY21.Chr11.1193 | KY21:KY21.Chr11.119 | 1261 | 60.98 | 0.47 | 2.6 | 0 | 129.74 | 23.45 | N.A. | complement factor H-like (LOC100180592), transcript variant X2, mRNA (XM_002125505.4) |
| KY21.Chr1.1023 | KY21:KY21.Chr1.1023 | 1236 | 60.26 | 1.85 | 0.05 | 0.05 | 32.57 | 1205.20 | 1205.20 | chromatin assembly factor 1 subunit A-A (LOC100175940), transcript variant X2, mRNA (XM_002120009.4) |
| KY21.Chr7.385 | KY21:KY21.Chr7.385.v | 707 | 59.82 | 0.17 | 0.1 | 0.1 | 351.88 | 598.20 | 598.20 | lens fiber membrane intrinsic protein-like (LOC100184210), mRNA (XM_002125020.2) |
| KY21.Chr2.183 | KY21:KY21.Chr2.183.v | 1486 | 59.41 | 0 | 1.63 | 0 | N.A. | 36.45 | N.A. | uromodulin-like (LOC100184343), transcript variant X2, mRNA (XM_026840819.1) |
| KY21.Chr1.406 | KY21:KY21.Chr1.406.v | 1425 | 59.1 | 0.88 | 0.04 | 0 | 67.16 | 1477.50 | N.A. | probable beta-tubulin polyglutamylase (LOC100177568), mRNA (XM_018815847.2) |
| KY21.Chr7.296 | KY21:KY21.Chr7.296.v | 1277 | 58.71 | 0.08 | 0.7 | 0 | 733.88 | 83.87 | N.A. | stabilin-2-like (LOC100186726), mRNA (XM_009860622.2) |
